# Conserved domains of meiosis-specific CHK2 protein in budding yeast contribute to its kinase activity

**DOI:** 10.1101/2023.04.01.535230

**Authors:** Jinsha Padmarajan, Abhijna Krishnan Edilyam, Vijayalakshmi V. Subramanian

## Abstract

Programmed double strand DNA breaks in meiosis can be repaired as inter-homologue crossovers and thereby aid the faithful segregation of homologous chromosomes. Biased repair mechanisms enforce repair with the homologue. Further, DNA breaks left unrepaired lead to checkpoint activation. Meiosis-specific Chk2 kinase in budding yeast mediates the biased repair of meiotic DSBs using homologue partner but also enforces the meiotic checkpoint. Here we investigate Mek1 kinase activity in budding yeast by analyzing novel point mutants derived from an EMS mutagenesis screen. The point mutants in different domains of Mek1 abolish its activity that cannot be rescued by complementation in transheterozygotes. Our findings lend insight on the mechanism of Mek1 function during meiosis.

## Introduction

Meiosis is a highly organized chromosome segregation process in sexually-reproducing organisms that generates haploid gametes from diploid progenitors (Kleckner 1996; Hochwagen 2008). Reduction in ploidy is achieved by a single round of chromosome duplication event followed by segregation of homologous chromosomes in meiosis I and that of sister chromatids in meiosis II (Kleckner 1996; Hunter 2015). Accurate homologue segregation in meiosis I entails induction of programmed double-stranded DNA breaks (DSBs) and subsequent repair as crossovers between the homologues (Subramanian and Hochwagen 2014; Lam and Keeney 2014; Hunter 2015). At least one crossover per homologue pair is required to create a physical link between them (Hochwagen 2008). This promotes proper orientation and segregation of the homologues on the meiotic spindle and preserves fertility during this process (Kleckner 1996; Hochwagen 2008).

While meiotic DSBs can potentially use either the homologue or the proximal sister chromatid as a template for repair, only repair from the homologue can give rise to crossovers that are crucial for accurate chromosome segregation (Subramanian and Hochwagen 2014; Hunter 2015). Therefore, unlike repair of DSBs in vegetative cells which primarily utilize the sister chromatid as a template (Kadyk and Hartwell 1992; Bzymek *et al*. 2010), DSBs in meiosis preferentially repair using the homologue as a template (Hollingsworth 2010; Humphryes and Hochwagen 2014). This biased repair with the homologue template is mechanistically linked to DSB repair in meiosis (Subramanian and Hochwagen 2014). In *S. cerevisiae*, DSB ends and the 5′ end-resected single stranded DNA ends trigger conserved checkpoint kinases Tel1/Mec1 (ATM/ATR orthologues), which set the stage for downstream repair of DSBs and also promote homologue bias (Grushcow *et al*. 1999; Cheng *et al*. 2013; Penedos *et al*. 2015). Additionally repair of meiotic DSBs occurs in the context of specialized meiotic chromosomal proteins which includes meiotic cohesins, Red1, Hop1 (conserved HORMAD protein) and Mek1 kinase (Chk2 orthologue) (Hollingsworth and Byers 1989; Rockmill and Roeder 1991; Leem and Ogawa 1992; Smith and Roeder 1997; Bailis and Roeder 1998; Klein *et al*. 1999; Panizza *et al*. 2011). The checkpoint kinases (Tel1/Mec1) phosphorylate Hop1 at threonine 318 in response to DSBs (Carballo *et al*. 2008; Cheng *et al*. 2013). Phosphorylated Hop1 serves as an adaptor for Mek1 kinase localization to meiotic chromosomes (presumably at/near DSB sites), its dimerization and activation of the kinase function (Niu *et al*. 2005, 2007; Carballo *et al*. 2008; Ontoso *et al*. 2013; Subramanian *et al*. 2016).

Chk2 kinase contributes to repair choices and outcome as well as recombination checkpoint in several organisms (Subramanian and Hochwagen 2014). In budding yeast, Mek1 kinase activity is responsible for promoting homologue bias; in *mek1Δ* or upon Mek1 kinase activity inhibition, the DSBs are repaired with the sister chromatid template and result in meiosis I segregation errors (Wan *et al*. 2004; Niu *et al*. 2007). While Mek1 restricts Rad51 strand invasion activity by phosphorylating its cofactor Rad54 and interactor Hed1, other Mek1-dependent mechanisms are also at play to promote homologue bias (Liu *et al*. 2014; Callender *et al*. 2016). Additionally, Mek1 contributes to crossover interference (a mechanism to ensure non-random and well-spaced crossover placement) by targeting synaptonemal complex proteins (Chen *et al*. 2015). Similarly, a reduction in homologue crossovers is also observed in *C. elegans* Chk2 orthologues, *cds1/2* mutants (Oishi *et al*. 2001). Further, Mek1/Chk2 facilitates meiotic recombination checkpoint and delays cell cycle progression when DSBs are unrepaired (Pérez-Hidalgo *et al*. 2008; Wu *et al*. 2010). In mouse, Chk2 is responsible for eliciting DNA damage response (Bolcun-Filas *et al*. 2014; Pacheco *et al*. 2015; Martínez-Marchal *et al*. 2020). One mechanism by which Mek1 elicits the recombination checkpoint in budding yeast is by phosphorylating and restricting the activity of Ndt80, a transcription factor that is essential for exit from meiotic prophase (Prugar *et al*. 2017; Chen *et al*. 2018).

Mek1 protein has three conserved domains with respect to Chk2; an N-terminal forkhead-association domain (FHA), a kinase domain and a C-terminal domain. Upon DSB formation, the FHA, a conserved phospho-protein binding domain of Mek1 binds to phosphorylated Hop1 (pHop1) (Durocher and Jackson 2002; Wan *et al*. 2004). Recruitment of Mek1 by pHop1 promotes its dimerization and autophosphorylation of threonine residues in its kinase activation loop, leading to kinase activation (Wan *et al*. 2004; Niu *et al*. 2007). Specific mutations in these domains have been reported to abolish the kinase activity of Mek1 (Wan *et al*. 2004; Niu *et al*. 2007).

Here we report analysis of novel *mek1* point mutations isolated from an EMS mutagenesis screen that either carry missense mutations in the FHA or kinase domains or truncate the C-terminus. We assess Mek1 function and protein stability in these mutants. We find that in the missense mutants the protein is stably expressed and the predicted structure of the mutant protein is unchanged. On the other hand, C-terminus truncated Mek1 mutants have an altered predicted structure and accordingly exhibit reduced protein levels in meiosis. Regardless, function is severely compromised in all mutants. Finally, genetic trans-complementation of these mutants suggest that all domains are essential even *in trans* in a dimer. Our findings indicate that the conserved domains of Mek1 are key to its kinase activity and for upholding homologue bias and recombination checkpoint in meiosis.

## Results and Discussion

Mek1 triggers a recombination checkpoint arrest when a meiosis specific strand invasion protein, Dmc1, is absent (Roeder and Bailis 2000; Wu *et al*. 2010). However if *MEK1* is deleted, the block on Rad51 strand invasion protein is lifted, DSBs are repaired (albeit with sister chromatids in the absence of homologue bias) and budding yeast cells complete meiosis to form gametes (spores) (Wan *et al*. 2004). Taking advantage of this mechanism, an EMS mutagenesis screen was performed to isolate mutants that bypass *dmc1* meiotic arrest (the screen will be described elsewhere). Point mutations in *mek1\** were isolated from the screen and identified using genetic non-complementation with *mek1Δ* followed by sanger sequencing. Here we discuss five mutations in the three conserved domains of Mek1 (Figure 1a); one in FHA domain (S66F), two in the kinase domain (P245L, P384S) and two at the C-terminus (W410*, W443*).

**Figure 1:**
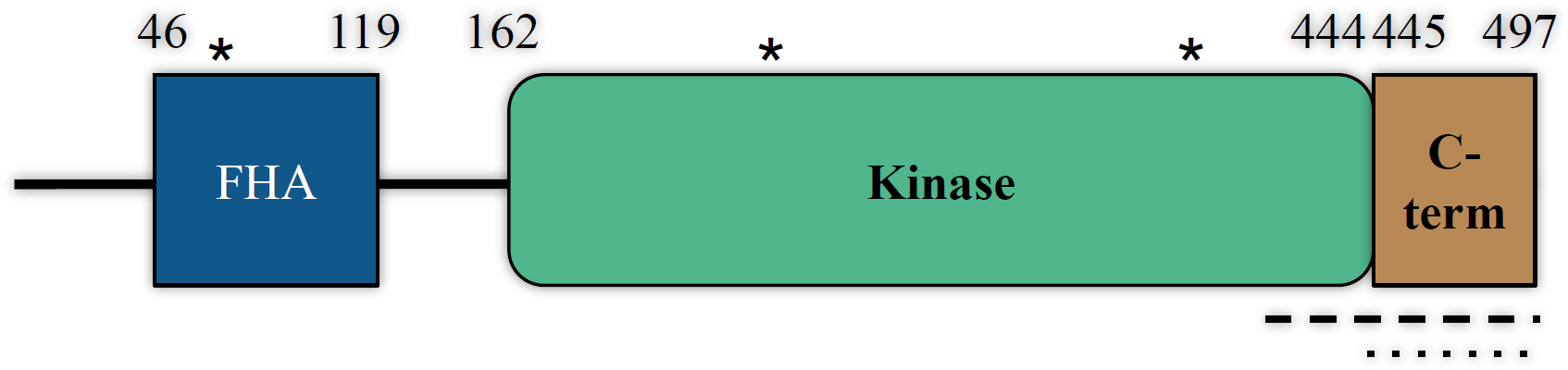
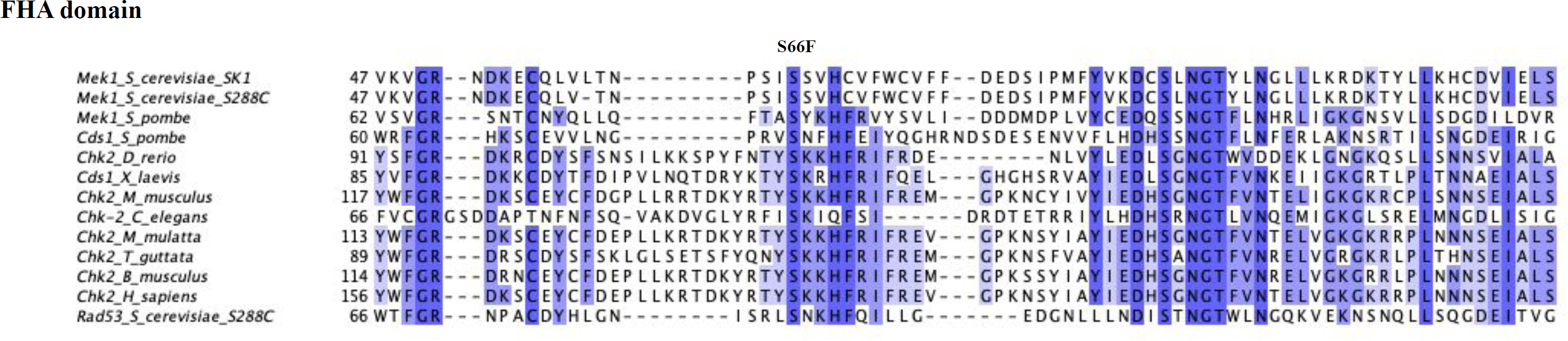
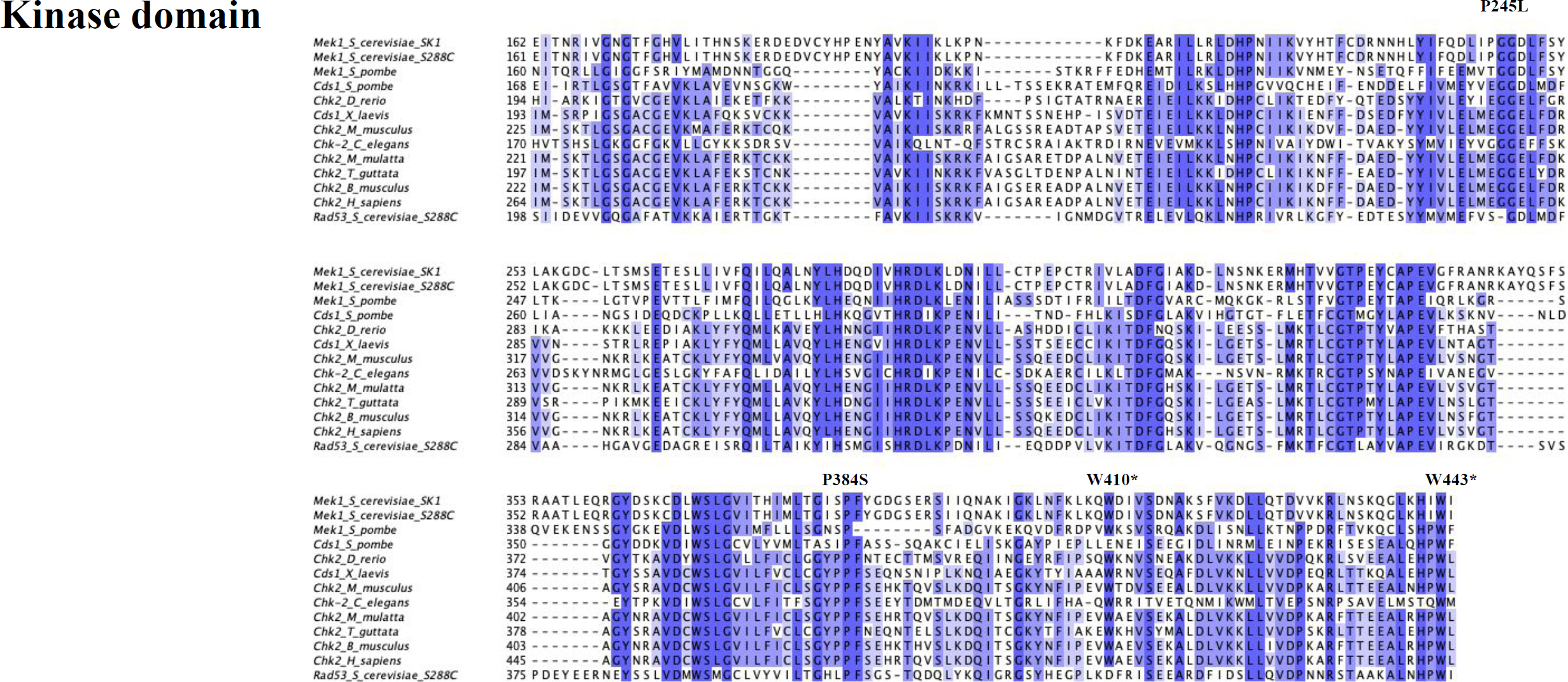
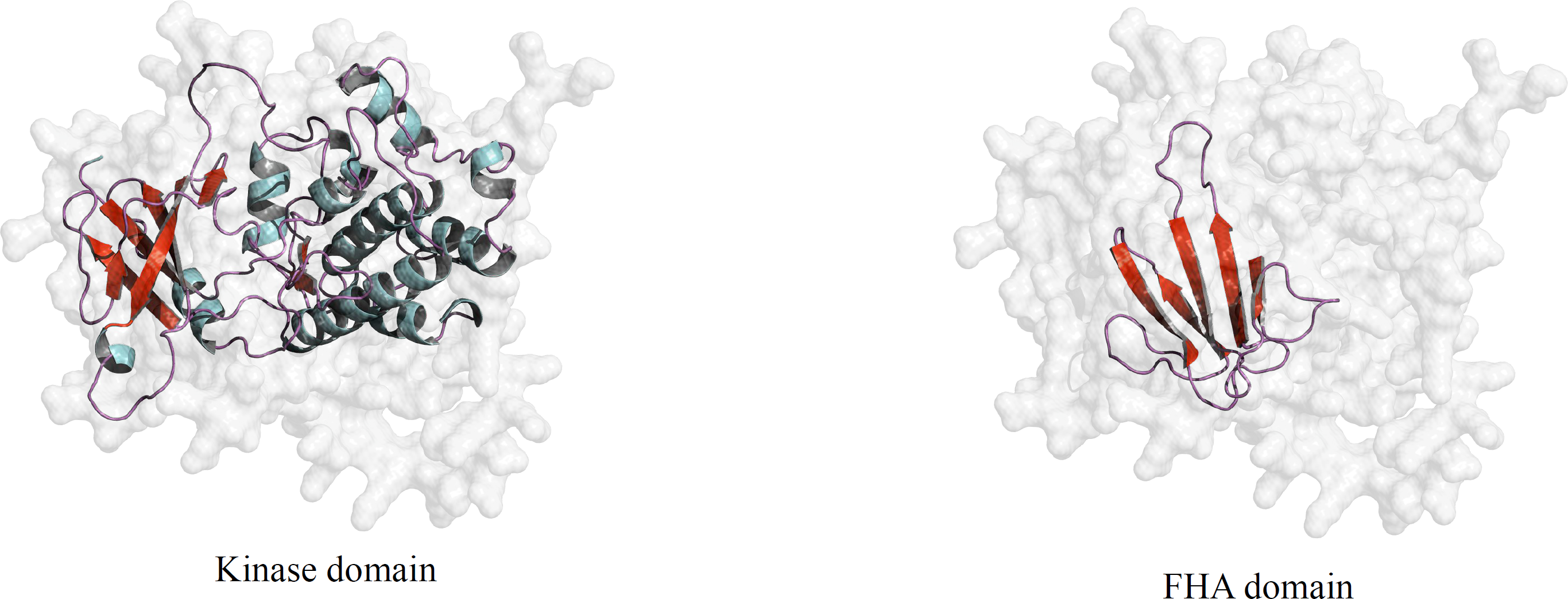
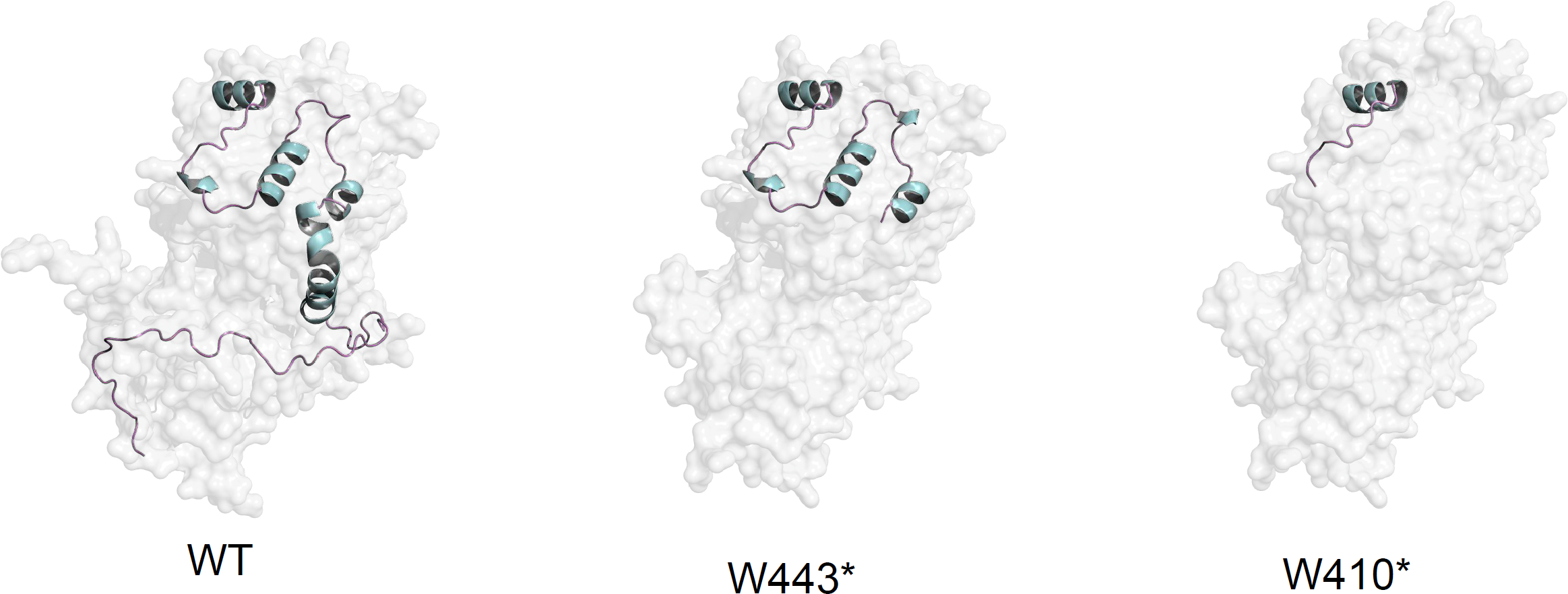
Sequence conservation and structure. a) Mek1 consists of FHA domain, catalytically active kinase domain and a conserved C-terminal domain; ⁎ denotes position of point mutations (S66F, P245L, P384S), dashed and dotted lines denotes W410* and W443* truncations respectively. b) Multiple sequence alignment of FHA domain; the position of the point mutation S66F is indicated c) Multiple sequence alignment of the kinase domain; the position of the point mutations (P245L, P384S) and truncations (W410*, W443*) are indicated. e) Alphafold2 predicted structures of kinase domain and FHA domain. The models are coloured based on secondary structures. f) Alphafold2 predicted C-terminal regions of wild type, Mek1-W443*, Mek1-W410*. C-terminal regions are highlighted as helices connected by loops using pymol. Mek1 C-terminal truncation mutants lack the C-terminal alpha helix and a disordered region compared to wild type.

### Mek1 kinase sequence and domains are conserved

Mek1 is a meiosis specific checkpoint kinase with three structural domains, namely FHA domain, kinase domain and C-terminal domain (Figure 1a) (Niu *et al*. 2007). Both the FHA and kinase domains are conserved across the Chk2 kinase family. A multiple sequence alignment using ClustalW revealed that mutant sites S66 and P384 are conserved across species while P245 is beside a conserved site G246 (Figure 1b, 1c). Alphafold2 prediction of Mek1 structure showed that the kinase domain consists of alpha helices and beta strands, while 73 amino acids long FHA domain is made up of 6 beta strands (Figure 1d) (Jumper *et al*. 2021; Mirdita *et al*. 2021). Interpro protein domain prediction was done to visualize possible loss of kinase and FHA domains. Interestingly, neither the mutation in the FHA domain (S66F) nor the two missense kinase mutations P245L and P384S dramatically affect the predicted structure of Mek1 (data not shown) suggesting that protein folding may not be the cause of the mutant phenotype. The C-terminal truncation mutants W410* and W443* have a deletion of 87 and 54 amino acids respectively. Alphafold2 predicted loss of three C-terminal alpha helices and a disordered region in W410*, while loss of one C-terminal helix and disordered region in W443* (Figure 1e). These findings suggest that the C-terminal domain of Mek1 may contribute to proper folding of Mek1 kinase.

### *mek1\** point mutants phenocopy *mek1Δ* mutant

The *mek1\** point mutants were isolated on the basis of their ability to bypass meiotic arrest observed in *dmc1Δ*, however, the strains may have several other mutations in the background. To clearly assign the phenotype to *mek1*, we introduced each point mutation into a non-mutagenized, clean yeast background using CRISPR Cas9. The *mek1\** point mutants were then assessed for their ability to bypass the recombination checkpoint in *dmc1Δ*. The point mutants rescued the sporulation defect seen in *dmc1Δ* (Figure 2a, Table S2a), suggesting that these point mutations in *mek1* abolished its recombination checkpoint activity. Additionally, none of the spores were viable (data not shown), suggesting a loss of homologue bias in the DSB repair process. We also assessed the phenotype of *mek1\** point mutants in the presence of Dmc1 strand exchange activity. No defects were observed in sporulation (Figure 2b, Tables S2b, S2c), however the spores were not viable (Table S2d). These findings indicate that *mek1\** point mutants phenocopy *mek1Δ* with respect to sporulation and spore viability and exhibit a complete loss of function.

**Figure 2:**
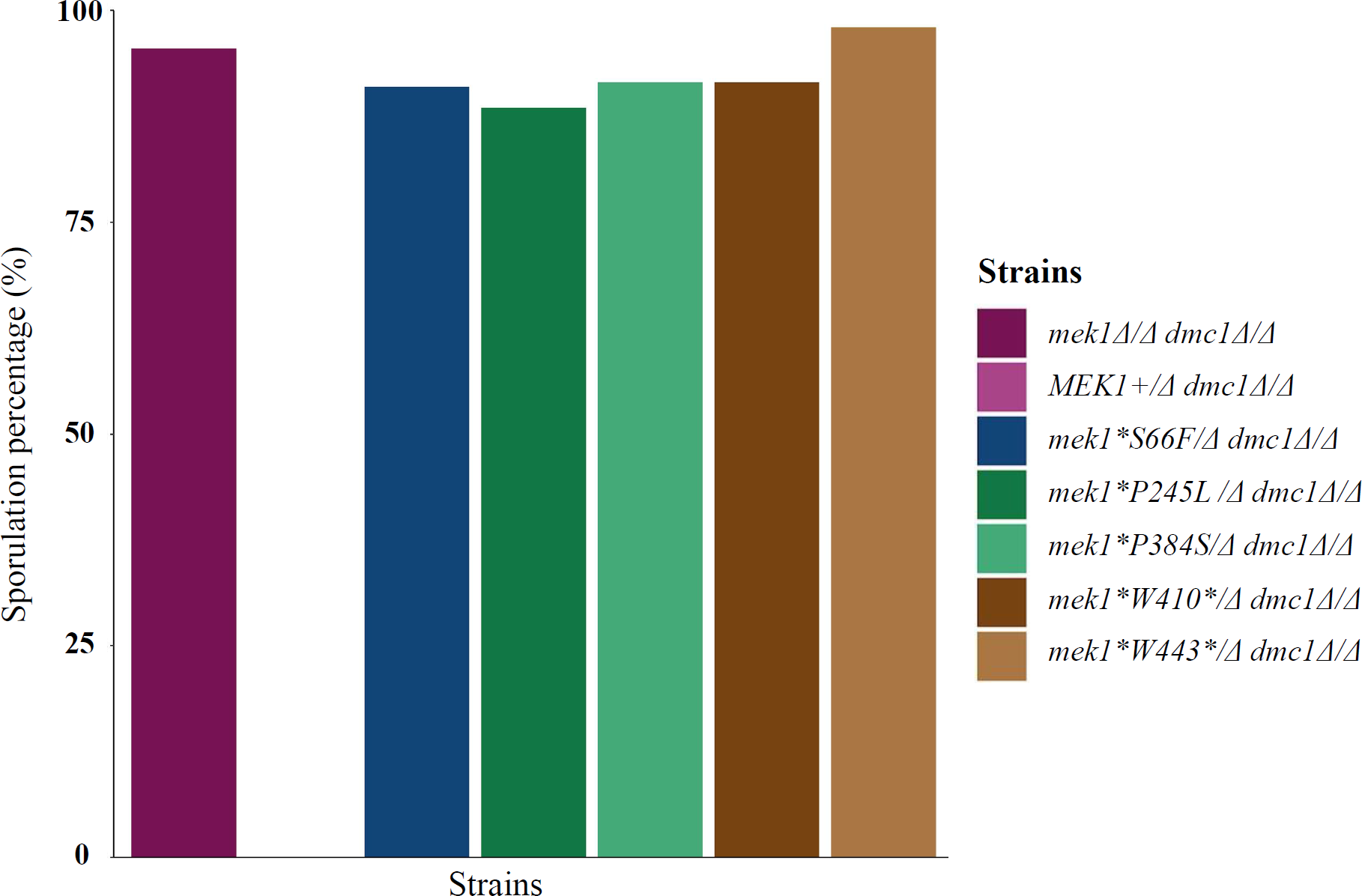
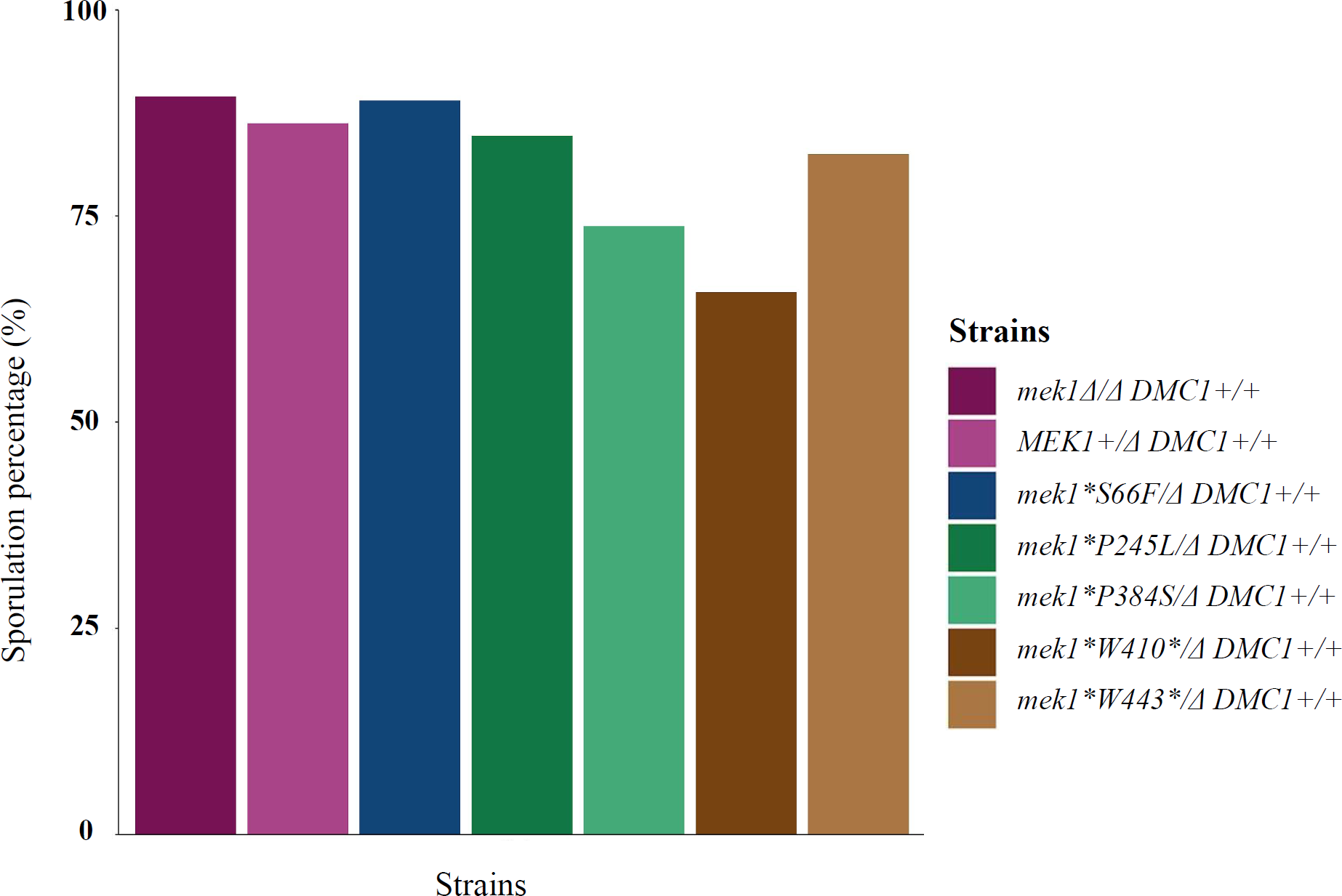
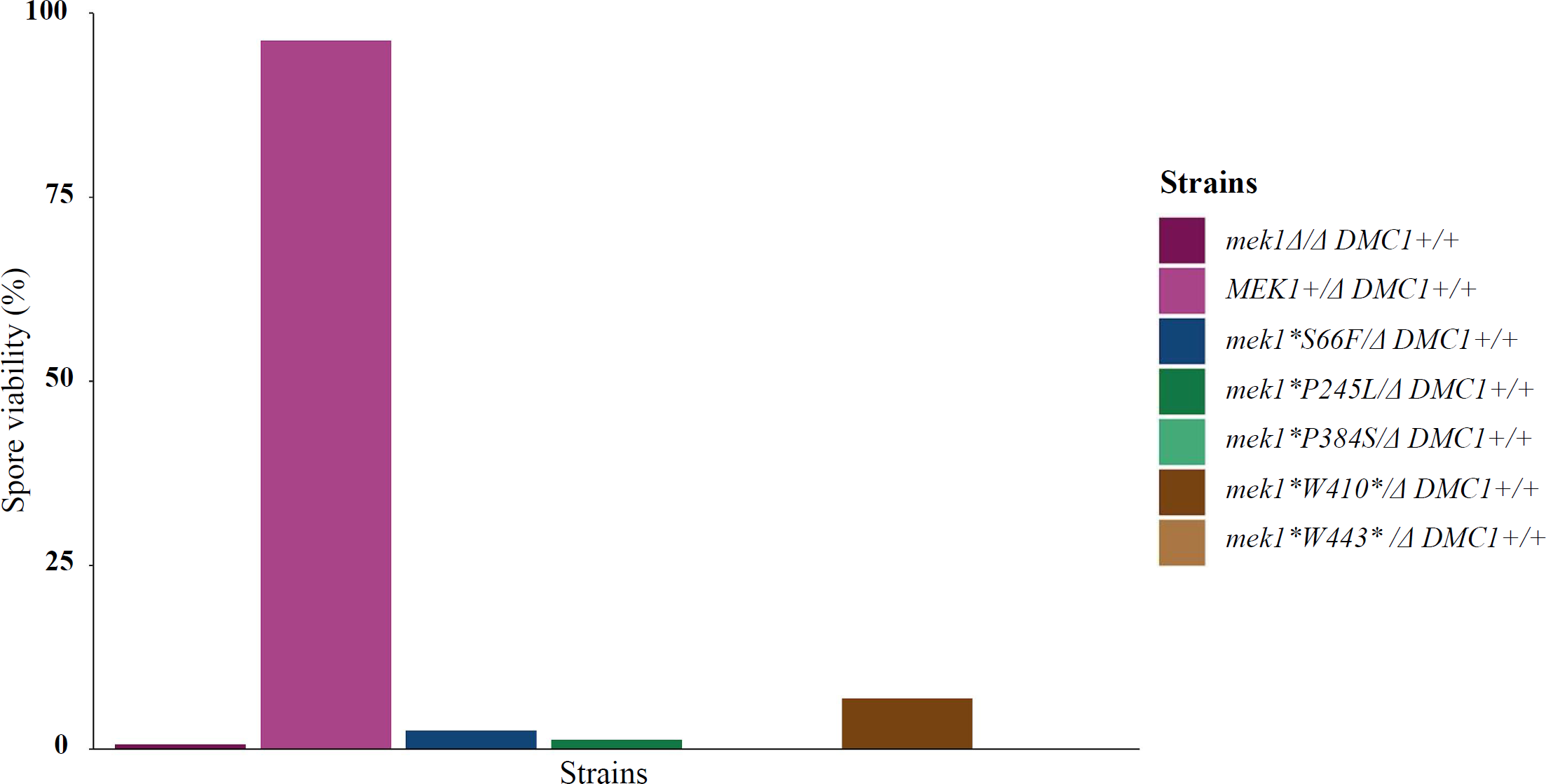
Sporulation efficiency and spore viability in *mek1* dmc1* and *mek1* DMC1* cells. a) Sporulation efficiency was defined as the percentage of the number of spores to the total number of cells counted. Sporulation efficiency of *mek1\** point mutants along with control in *dmc1* background was calculated for n=200 for each of the indicated strains. b) Sporulation efficiency of *mek1\** point mutants in DMC1 background was calculated as an average from 2 biological replicates for n=200 c) Spore viability was defined as the percentage ratio of number of spores that are viable divided by the total number of spores dissected.

### Mek1 point mutations do not dramatically affect protein stability

Because *mek1\** point mutants exhibited a complete loss of function and phenocopied *mek1Δ*, we wondered if protein folding and stability were compromised leading to loss of function. To interrogate the protein levels in the *mek1\** point mutants, we did a western analysis to assess levels of Mek1 protein during meiosis (Figure 3, Figure S1). We used meiosis-specific Red1 as well as Pgk1 as our loading controls. The Mek1 protein levels were similar in the missense mutants compared to wild type. These findings, in addition to absence of any predicted folding defects in these mutants, suggest that in the FHA and kinase domain mutants loss of function may be due to loss of critical interactors rather than the protein itself. The C-terminal truncation mutants, however, exhibited lower Mek1 protein levels (Figure 3, Figure S1). The predicted folding of the C-terminal truncation is also compromised. Together these observations suggest that the truncation mutants might be less stable.

**Figure 3:**
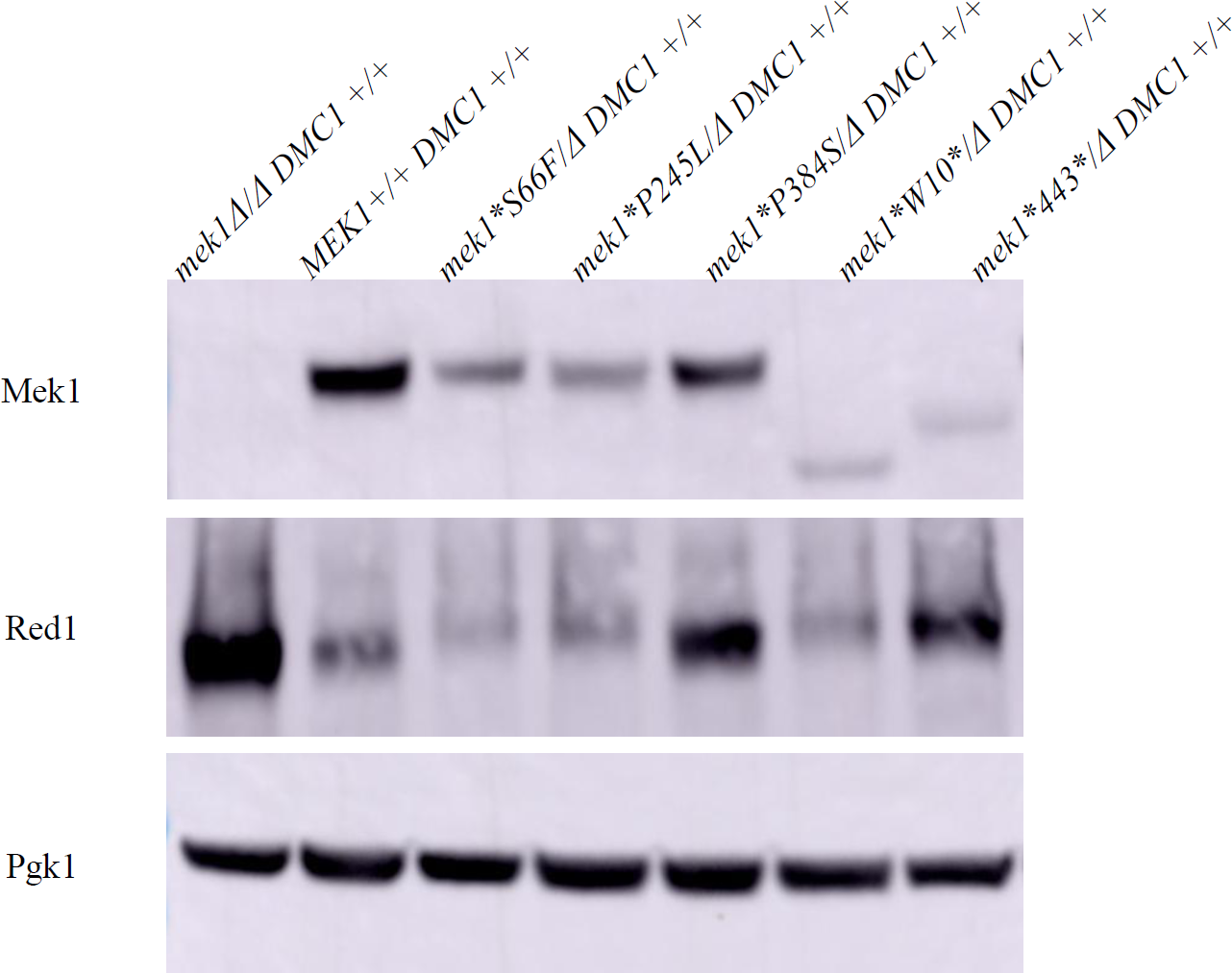
Mek1 protein is stably expressed in the point mutant strains. Western blot analysis showing Mek1 protein expression for the indicated *mek1\** point mutants in a *DMC1* background. Samples were taken at 4

### Kinase activity of Mek1 point mutants is severely compromised

In FHA and kinase mutants, the protein levels were not compromised, however, these mutants phenocopied *mek1Δ*. To identify the cause of phenotypic defects in these mutants we sought to assess their kinase activity. Substrates of Mek1 include phosphorylation of histone H3 on threonine 11 (Govin *et al*. 2010; Subramanian *et al*. 2016; Kniewel *et al*. 2017). Phosphorylation of H3T11 was not observed in the FHA and kinase mutants (Figure 4, Figure S2). This lack of Mek1 activation was not because DSB formation was compromised; γ-H2AX signal (a marker downstream of DSBs) was similar to controls (Figure 4, Figure S2). Very low levels of pH3T11 signal was observed in C-terminal truncation (W443*, see Figure S2) also pointing to why these cells exhibited mutant phenotype. These findings are consistent with previous reports (Wan *et al*. 2004; Niu *et al*. 2007), where point mutations in the FHA domain or the C-terminal regions caused loss of Mek1 activity.

**Figure 4a:**
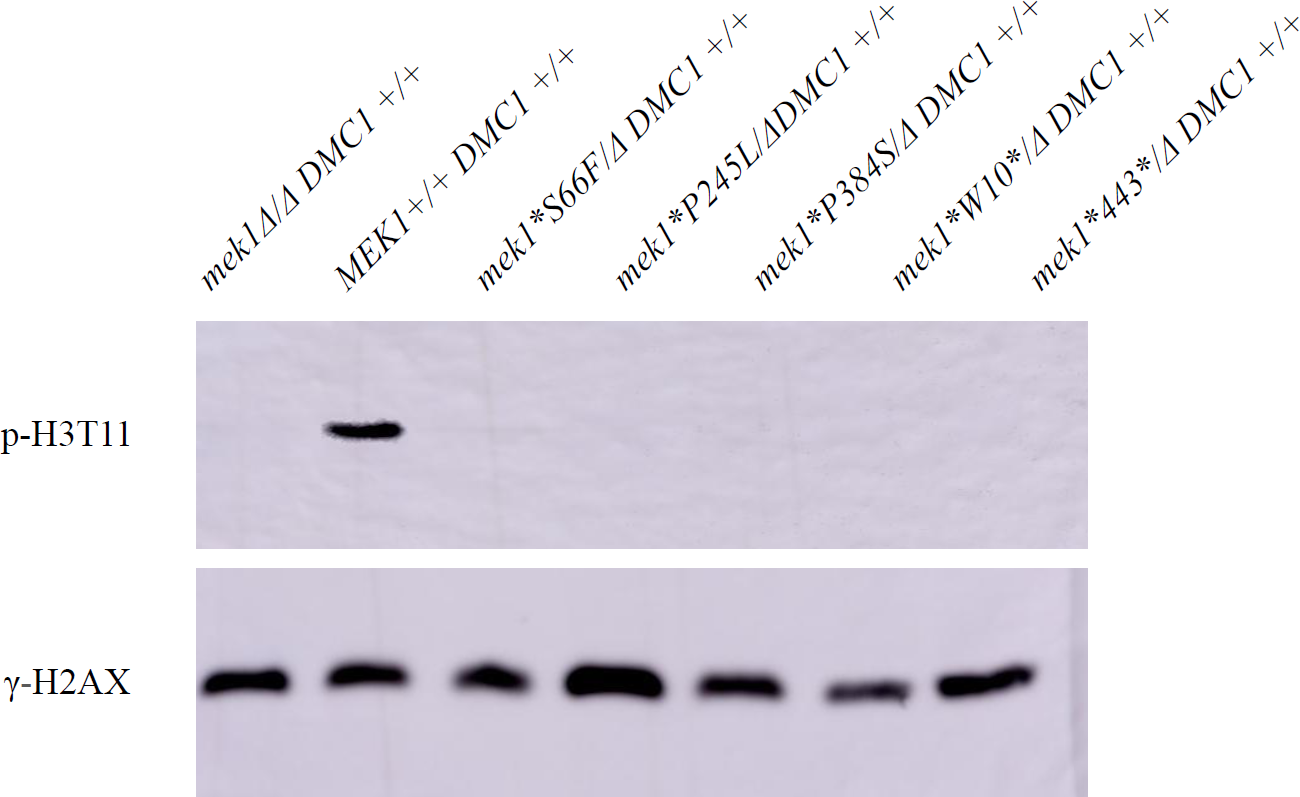
Mek1 kinase activity is not observed in the point mutants. Mek1 kinase activity was monitored by the phosphorylation of Histone 3 at threonine 11. Cells harvested at 4 hour time point were resolved by SDS-PAGE and analyzed using western blot by probing with anti-phospho-H3T11 antibody for the indicated diploid strains (upper panel). Phosphorylation of γ-H2AX at Serine 129 was used as a marker for DSB formation (lower panel).

### Transheterozygotes of *mek1* point mutants do not rescue the mutant phenotype

None of the *mek1\** mutants analyzed so far were functionally competent. However, because the Mek1 function *in vivo* requires its dimerization, one possibility is that the mutations in different domains may complement each other *in trans* following dimerization. To examine complementation between different alleles *in trans*, we generated *dmc1Δ* transheterozygotes. We did observed a rescue of *dmc1Δ* meiotic arrest suggesting that each of these domains are critical for Mek1 function even within a dimer. It is also possible that these domains are important for dimerization itself. Indeed ectopic dimerization rescued function of point mutations in FHA and C-terminus of Mek1 (Wan *et al*. 2004; Niu *et al*. 2007).

Our work has investigated novel *mek1* mutants. We have found that each of the Mek1 domains are critical for its function. The mutants are unable to impose recombination checkpoint or homologue bias. Interestingly the mutants are also able to rescue the *dmc1* arrest *in trans* indicative of lack of Mek1 activity. Further analysis of these mutants will shed light on the mechanisms of Mek1-mediated homologue bias and checkpoint.

## Materials and methods

### Strains

All experiments were performed using strains in the SK1 background. The list of strains can be found in Table S1. *mek1\** point mutant strains were derived from a random EMS (Ethyl methanesulfonate) mutagenesis of *dmc1* mutants. The *mek1\** gene amplicon of the mutant alleles were transformed into a non-mutagenized genetic background (*mek1Δ::kanMX*) along with CRISPR Cas9 plasmid also carrying guide RNA targeting *KanMX* (pRS425-*Cas9-sg-KanMX*). The cells were plated on leucine-dropout (DO) selection media to select for cells transformed with the Cas9 plasmid which would cleave at the *KanMX* sequence. The obtained transformants were streaked on YPD media to lose the plasmid. *mek1\** point mutants clones for further study clones were chosen on the basis of negative selection on both leucine-DO and G418 drug plates. The *mek1\** point mutants were mated with a *mek1* null strain in *dmc1* as well as *DMC1* background and the obtained diploids were used for functional assays.

### Structure and functional domain prediction and alignment of Mek1

Structure prediction of proteins was done using Deepmind’s Alphafold2, using the open access alphafold2_advanced script (https://colab.research.google.com/github/sokrypton/ColabFold/blob/main/beta/AlphaFold2_advanced.ipynb)(Jumper *et al*. 2021; Mirdita *et al*. 2021). Mmseq2 was used as multiple alignment tool for all the predictions with rest of the parameters as default. This method was used to generate five predicted models with output as pdb file. The ranks were assigned based on the pLDDT score, and the model with the highest rank was used for further analysis. The generated alpha fold structures were visualised using pymol (Schrodinger, LLC. 2010. The PyMOL Molecular Graphics System, Version 1.20). Protein domain prediction of mutants was done using EBI’s InterPro (https://www.ebi.ac.uk/interpro/). Multiple sequence alignment was done online using the MAFFT tool from EBI website (https://www.ebi.ac.uk/Tools/msa/mafft/) and visualised using Jalview(Waterhouse *et al*. 2009). Sequences were sorted based on pairwise identity with Mek1 from *S cerevisiae* (SK1) as reference sequence.

### Synchronous meiosis

Freshly patched diploid strains on YPG were inoculated in YPD culture and grown at room temperature while shaking at 200rpm for 24hrs. Cells were diluted in 50ml BYTA (50mM sodium phthalate-buffered, 1% yeast extract, 2% tryptone and 1% acetate) to a final A_600_ =0.3 and incubated for 16hrs at 30°C at 200rpm. Meiosis was induced in sporulation media (0.3% potassium acetate, 5% acetic acid) by diluting the cells to an O.D of 2. Culture was incubated at 30°C at 200rpm and samples were collected at desired time points.

### Sporulation and spore viability assays

Sporulation of the diploid strains were monitored after 24hrs of meiotic induction. For each biological replicate, a total of 200 spores were counted and are shown as a percentage of sporulated cells from a population of cells to obtain sporulation efficiency. For spore viability, 20μl sporulated samples were added to equal volume of 1mg/ml Zymolyase T-100 and incubated at 37°C for 30 mins. Zymolase treated tetrads were dissected and the spores were allowed to germinate on growth media for two days. Spore viability was determined as the percentage of viables spore clones from total spores dissected. The graphs for sporulation efficiency and spore viability for the the diploid mutants strains along with controls were plotted using R version 4.2.2.

### Whole-cell protein extraction and Western blot analysis

For protein extraction, 5ml of samples were collected from the sporulating culture at required time intervals. The cell pellets were resuspended in 5ml of 5% trichloro acetic acid (TCA). Cells were then washed in 1M unbuffered Tris and resuspended in TE (50mM Tris pH 7.5, 1mM EDTA) + DTT (200mM final concentration) master mix. Samples were boiled at 95°C for 5 min in SDS sample buffer and stored at -20°C for long term storage. Protein samples were resolved on SDS-PAGE followed by transfer onto nitrocellulose membrane at a constant voltage of 100V for 1hr. Mek1 and Red1 proteins were resolved on an 8% acrylamide resolving gel whereas phosphorylated histones were resolved on a 15% acrylamide gel.

### Antibodies

The details of primary and secondary antibodies used in this study are listed in supplementary table S3.

### Trans-complementation with *mek1\** mutants

Patches of each *mek1\** point mutant in a *MATa dmc1 ade2 LEU2* background was replica-plated onto a lawn of each *mek1\** point mutant in a *MATa dmc1 ade2 LEU2*. Diploids were selected on adenine-leucine-DO media. The diploids thus formed were transferred onto Protran BA85 nitrocellulose blotting membrane on adenine-leucine-DO selective media by replica plating. The plates were incubated at 30°C until a good amount of cells were grown on the membrane. The membranes were then transferred onto MSM plates (2% KOAc/AcAcid) to induce sporulation at 30°C for 4-5 days. The plates were screened under short UV for the presence of fluorescent patches (spores have a di-tyrosine component that fluoresces under UV light).

## Acknowledgements

We thank Andreas Hochwagen for generously sharing *mek1* mutants from the EMS mutagenesis screen. This work was supported in part by Ramalingaswami re-entry fellowship (BT/RLF/Re-entry/04/2019), DBT-Wellcome Trust India Alliance intermediate fellowship (IA/I/21/1/505647), SERB CRG grant (CRG/2021/007107/BHS) to V.V.S., and CSIR-JRF fellowship (09/1178(12250)/2021-EMR-I) to A.K.E.

---

**Table 1:** Trans-complementation between mek1* point mutants.

| **Mek1**<br>**genotype** | ***mek1Δ*** | ***MEK1+*** | ***mek1\*S66F*** | ***mek1\*P245L*** | ***mek1\*P384S*** | ***mek1\*W41***<br>***0\**** | ***mek1\****<br>***W443\**** |
| --- | --- | --- | --- | --- | --- | --- | --- |
| ***mek1Δ*** | S | NS | S | S | S | S | S |
| ***MEK1+*** | NS | NS | NS | NS | NS | NS | NS |
| ***mek1\*S66F*** | S | NS | S | S | S | S | S |
| ***mek1\*P245L*** | S | NS | S | S | S | S | S |
| ***mek1\*P384S*** | S | NS | S | S | S | S | S |
| ***mek1\*W410\**** | S | NS | S | S | S | S | S |
| ***mek1\*W443\**** | S | NS | S | S | S | S | S |

## Supplementary figures

**Figure S1:**
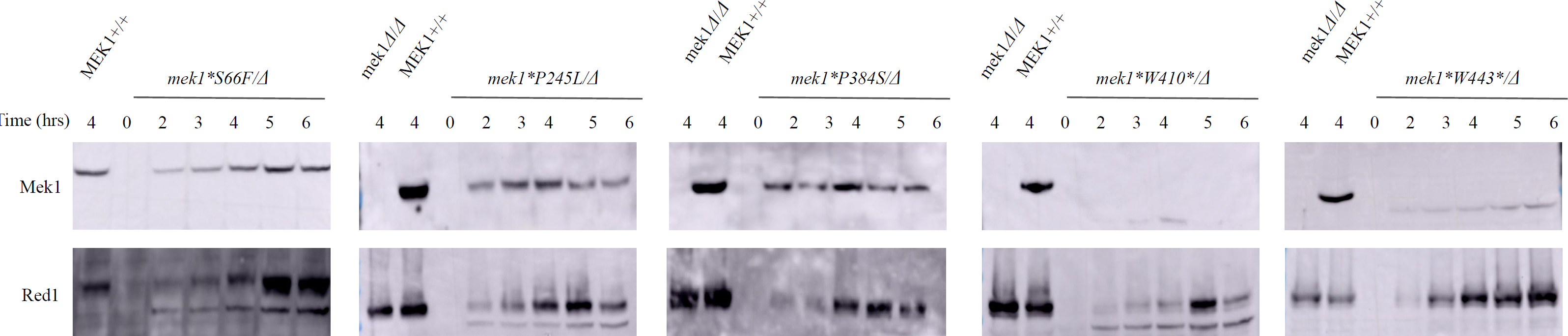
Mek1 expression during a time course in *mek1\** point mutants. Diploids of the indicated *mek1*/Δ* point mutants (DMC1 background) were induced to undergo meiosis in sporulation media and assayed at different time points to look at Mek1 protein expression by western blot. Red1 was used as a meiotic control. *mek1Δ/Δ* and MEK1+/+ was used as negative and positive controls respectively at 4 hour time point.

**Figure S2:**
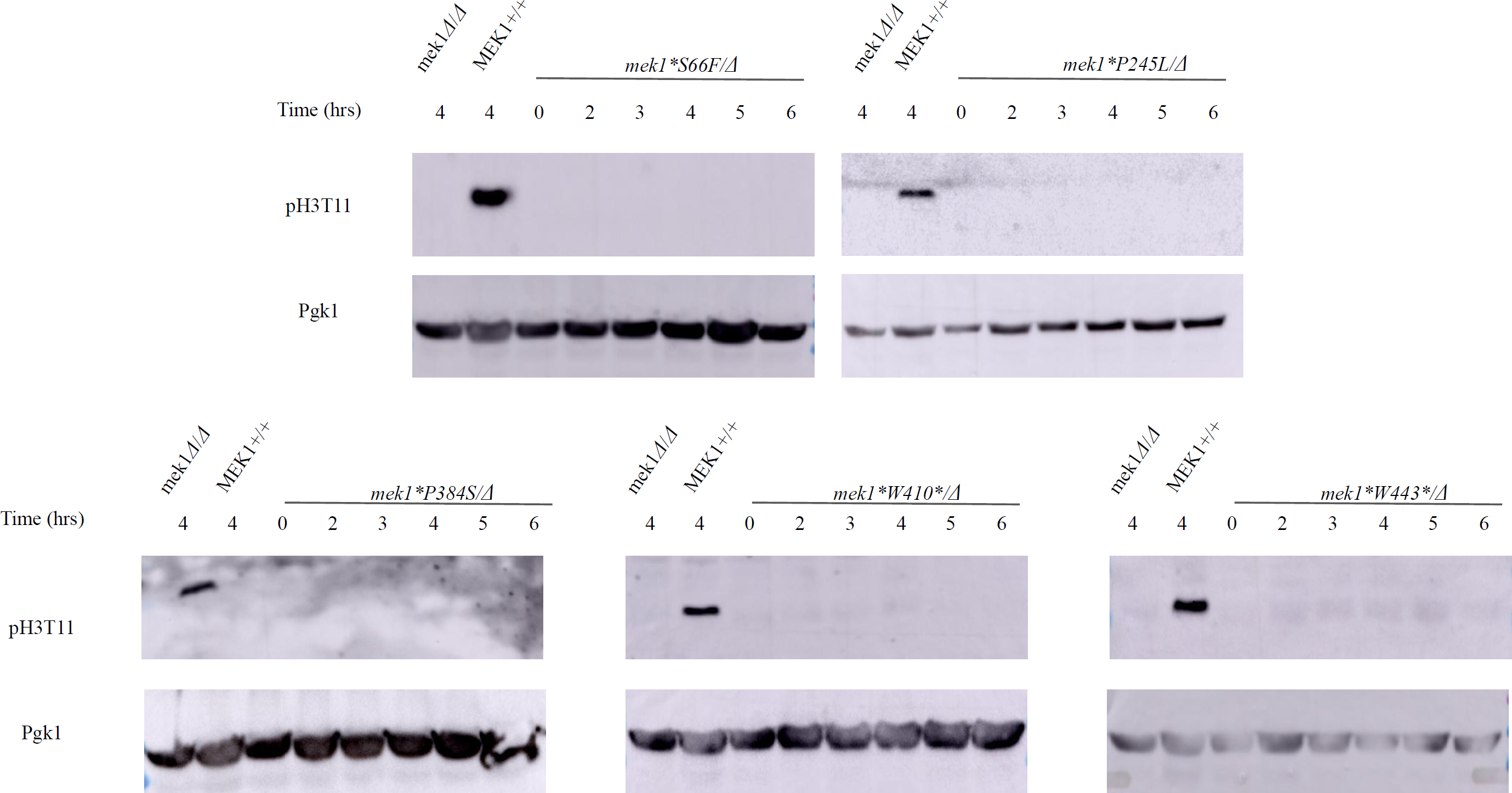
m*e*k1*\** point mutants are defective for H3T11 phosphorylation. Protein extracts for the indicated diploids were assayed by western blot at different time point to look for phosphorylation of Histone3 at T11 as a readout of kinase activity of *mek1\** point mutants (upper panel). Pgk1 was used as a loading control (lower panel).

**Table S1:**
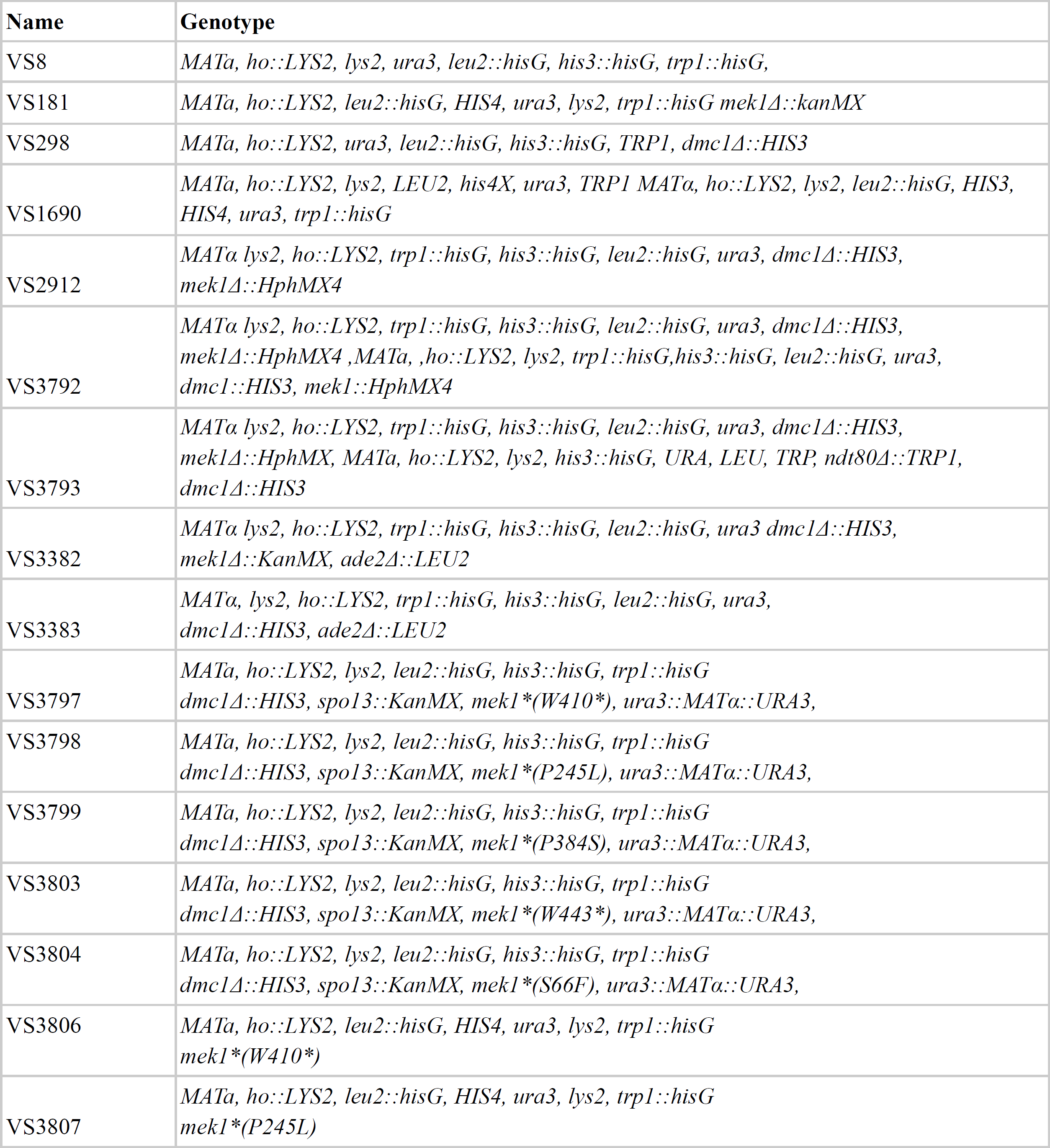
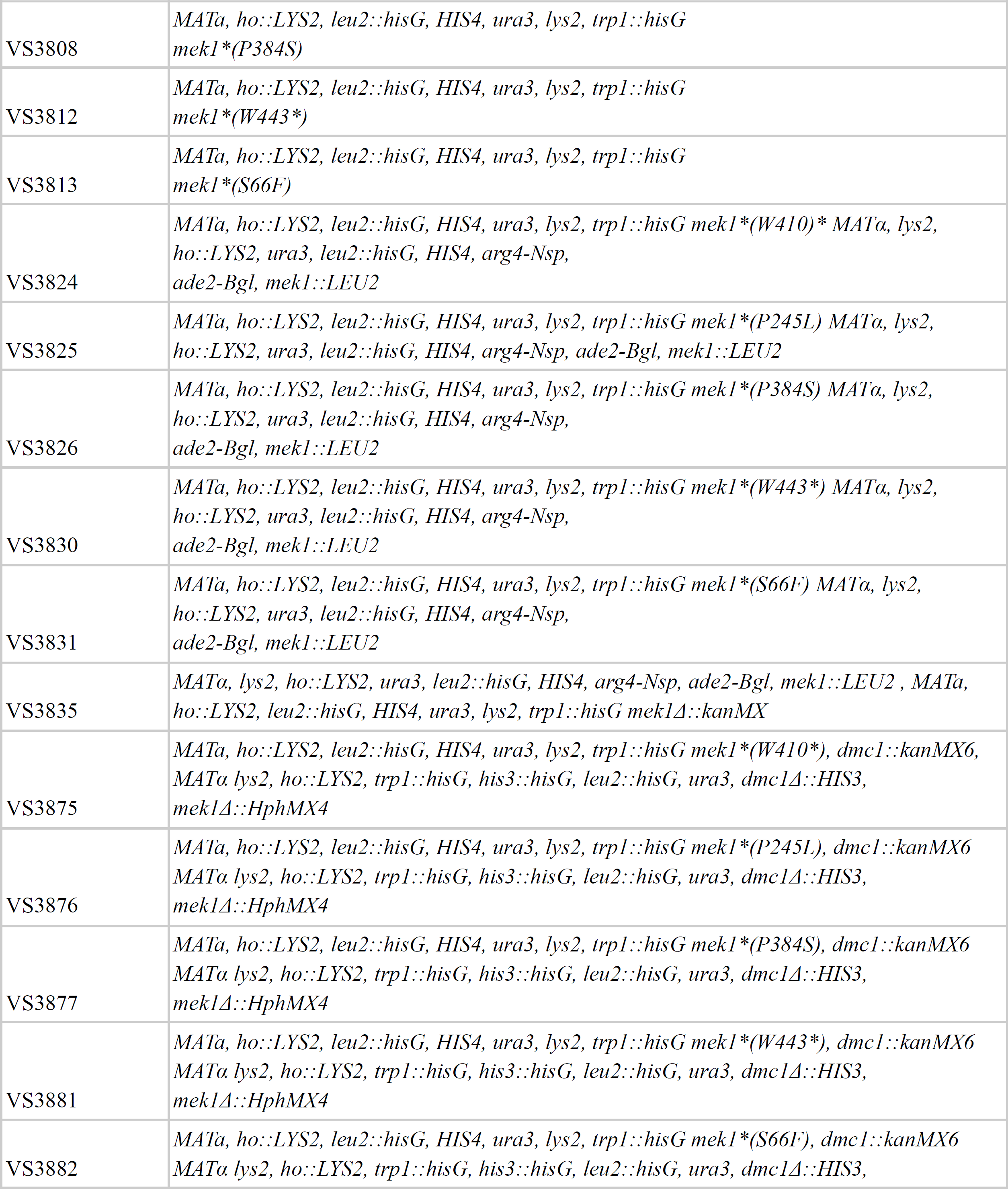
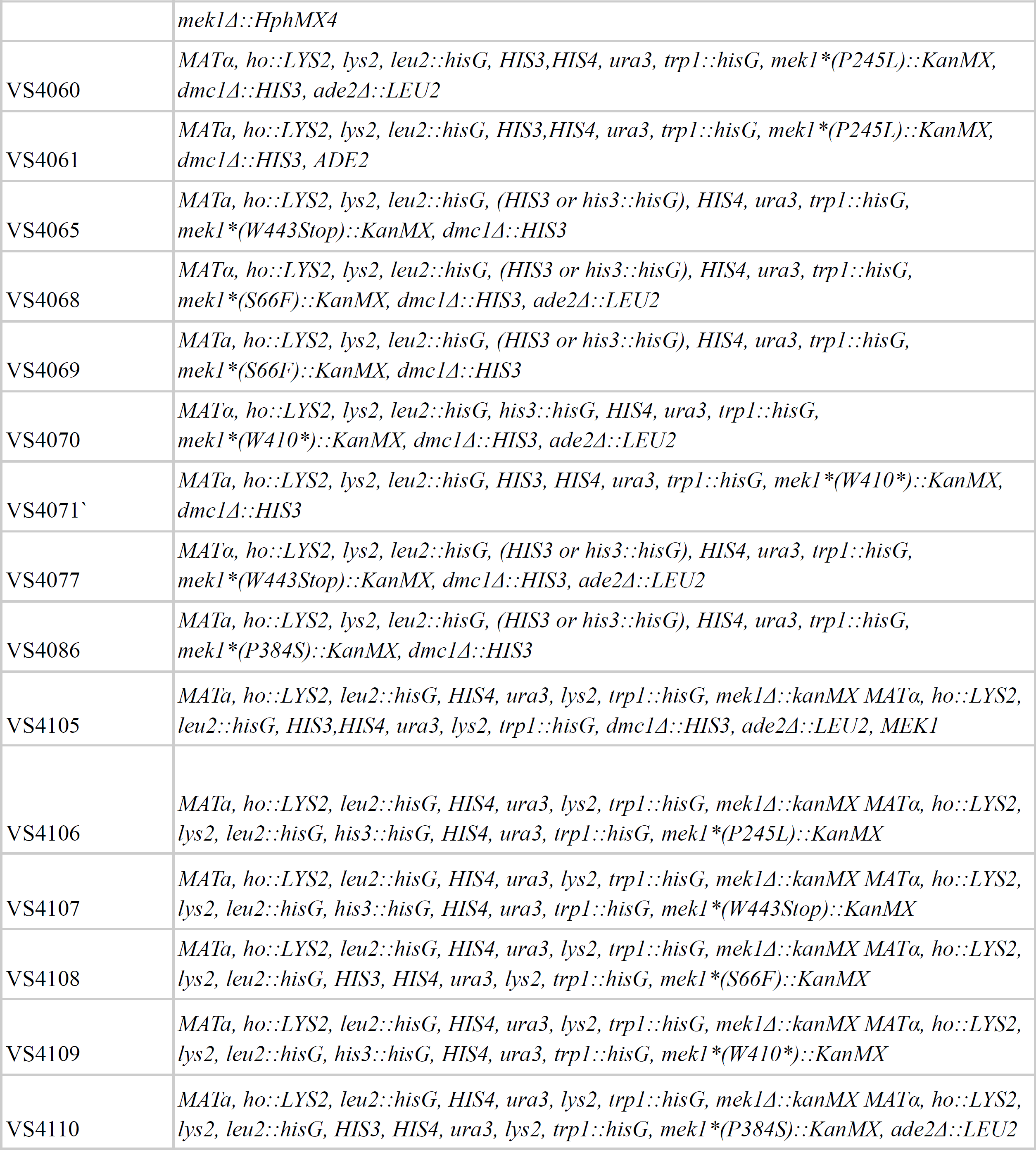
List of all *Saccharomyces cerevisiae* strains used in the study.

**Table S2a:** Sporulation efficiency of *mek1* point mutants in *dmc1* background.

| **Genotype** | **Strain**<br>**(#)** | **Unsporulated** | **Monads** | **Dyads** | **Tetrads** | **N** | **Sporulation (%)** |
| --- | --- | --- | --- | --- | --- | --- | --- |
| *mek1Δ/Δ dmc1Δ/Δ* | 3792 | 9 | 4 | 2 | 185 | 200 | 95 |
| *MEK1+/Δ dmc1Δ/Δ* | 3793 | 200 | 0 | 0 | 0 | 200 | 0 |
| *mek1\*S66F/Δ dmc1Δ/Δ* | 3882 | 18 | 1 | 25 | 156 | 200 | 91 |
| *mek1\*P245L/Δ dmc1Δ/Δ* | 3876 | 23 | 1 | 16 | 160 | 200 | 88.5 |
| *mek1\*P384S/Δ dmc1Δ/Δ* | 3877 | 17 | 2 | 13 | 168 | 200 | 91.5 |
| *mek1\*W410\*/Δ dmc1Δ/Δ* | 3875 | 17 | 1 | 22 | 160 | 200 | 91.5 |
| *mek1\*W443\*/Δ dmc1Δ/Δ* | 3881 | 4 | 1 | 32 | 163 | 200 | 98 |

**Table S2b:** Sporulation efficiency of mek1 mutants in DMC1 background (Set1)

| **Genotype** | **Strain (#)** | **Unsporulated** | **Monads** | **Dyads** | **Tetrads** | **N** | **Sporulation (%)** |
| --- | --- | --- | --- | --- | --- | --- | --- |
| *mek1*$\Delta/\Delta$ *DMC1*$+/+$ | 3835 | 18 | 3 | 6 | 163 | 200 | 86 |
| *MEK1*$+/+$ *DMC1*$+/+$ | 1690 | 12 | 1 | 2 | 185 | 200 | 94 |
| *mek1* * *S66F* $\Delta$ *DMC1* $+/+$ | 3831 | 5 | 3 | 7 | 185 | 200 | 97.5 |
| *mek1* * *245L* $\Delta$ *DMC1* $+/+$ | 3825 | 10 | 3 | 16 | 171 | 200 | 95 |
| *mek1* * *P384S* $\Delta$ *DMC1* $+/+$ | 3826 | 36 | 1 | 27 | 136 | 200 | 82 |
| *mek1* * *W410* $\Delta$ *DMC1* $+/+$ | 3824 | 16 | 1 | 16 | 167 | 200 | 92 |
| *mek1* * *W443* $\Delta$ *DMC1* $+/+$ | 3830 | 24 | 3 | 9 | 164 | 200 | 88 |

**Table S2c:** Sporulation efficiency of *mek1* mutants in *DMC1* background (Set2)

| **Genotype** | **Strain**<br>**(#)** | **No spores** | **Monads** | **Dyads** | **Tetrads** | **N** | **Sporulation**<br>**(%)** |
| --- | --- | --- | --- | --- | --- | --- | --- |
| *mek1*$\Delta/\Delta$ *DMC1*$+/+$ | 3835 | 14 | 4 | 5 | 177 | 200 | 93 |
| *MEK1*$+/+$*DMC1*$+/+$ | 4105 | 43 | 1 | 20 | 136 | 200 | 78.5 |
| *mek1* * *S66F* $\Delta$ *DMC1* $+/+$ | 4108 | 39 | 3 | 57 | 101 | 200 | 80.5 |
| *mek1* * *P245L* $\Delta$ *DMC1* $+/+$ | 4106 | 51 | 5 | 18 | 126 | 200 | 74.5 |
| *mek1* * *P384S* $\Delta$ *DMC1* $+/+$ | 4110 | 69 | 1 | 29 | 101 | 200 | 65.5 |
| *mek1* * *W410* $\Delta$ *DMC1* $+/+$ | 4109 | 121 | 0 | 8 | 71 | 200 | 39.5 |
| *mek1* * *W443* $\Delta$ *DMC1* $+/+$ | 4107 | 46 | 6 | 16 | 132 | 200 | 77 |

**Table S2d:** Spore viability of mek1 mutants in DMC1 background.

| Genotype | 4- viable | 3-viable | 2-viable | 1-viable | 0-viable | Total viable spore | N | Viability (%) |
| --- | --- | --- | --- | --- | --- | --- | --- | --- |
| *mek1Δ/Δ DMC1+/+* | 0 | 0 | 0 | 1 | 39 | 1 | 40 | 0.625 |
| *MEK1+/Δ DMC1+/+* | 35 | 4 | 1 | 0 | 0 | 40 | 40 | 96.25 |
| *mek1\*S66F/Δ DMC1+/+* | 0 | 0 | 1 | 2 | 38 | 3 | 40 | 2.5 |
| *mek1\*P245L/Δ DMC1+/+* | 0 | 0 | 0 | 2 | 38 | 2 | 40 | 1.25 |
| *mek1\*P384S/Δ DMC1+/+* | 0 | 0 | 0 | 0 | 0 | 0 | 40 | 0 |
| *mek1\*W410/Δ DMC1+/+* | 0 | 0 | 4 | 3 | 35 | 7 | 40 | 6.875 |
| *mek1\*W443/Δ DMC1+/+* | 0 | 0 | 0 | 0 | 40 | 0 | 40 | 0 |

**Table S3:** Antibodies used in the study.

| **Primary antibody** | **Dilution** | **Source** | **Secondary antibody** | **Dilution** | **Source** |
| --- | --- | --- | --- | --- | --- |
| anti-Mek1 | 1:10,000 | Pedro San-Segundo | anti-rabbit | 1:5000 | Abcam-Goat Anti-Rabbit IgG H&L (HRP) (ab6721) |
| ant-Red1 | 1: 5000 | Nancy Hollingsworth | anti-rabbit | 1:5000 | Abcam-Goat Anti-Rabbit IgG H&L (HRP) (ab6721) |
| anti-phospho-H3T11 | 1:1000 | Millipore (MC83) | anti-rabbit | 1:5000 | Abcam-Goat Anti-Rabbit IgG H&L (HRP) (ab6721) |
| anti-Pgk1 | 1:10,000 | Thermofisher (2C5D8) | anti-mouse | 1:5000 | Abcam-Goat Anti-mouse IgG H&L (HRP) (ab97019) |
| Anti-Histone H2A (phospho S129) | 1:1000 | Abcam (ab15083) | anti-rabbit | 1:5000 | Abcam-Goat Anti-Rabbit IgG H&L (HRP) (ab6721) |

